# Nonsense-mediated decay masks cryptic splicing events caused by TDP-43 loss

**DOI:** 10.1101/2025.07.09.664014

**Authors:** Yi Zeng, Odilia Sianto, Anastasiia Lovchykova, Chang Liu, Tetsuya Akiyama, Leonard Petrucelli, Aaron D. Gitler

## Abstract

In frontotemporal dementia and amyotrophic lateral sclerosis, the RNA-binding protein TDP-43 is lost from the nucleus, leading to cryptic exon inclusion events in dozens of neuronal genes. Here, we show that many cryptic splicing events have been missed by standard RNA-sequencing analyses because they are substrates for nonsense-mediated decay. By inhibiting nonsense-mediated decay in neurons we unmask hundreds of novel cryptic splicing events caused by TDP-43 depletion, providing a new picture to TDP-43 loss of function in neurons.

## Main

Amyotrophic lateral sclerosis (ALS) and frontotemporal dementia (FTD) are devastating neurodegenerative diseases with overlapping clinical and pathological features^1^. A hallmark shared by the majority of ALS and nearly half of FTD cases is the abnormal aggregation of the RNA-binding protein TDP-43^2^. TDP-43 pathology includes its mislocalization from the nucleus to the cytoplasm, accompanied by the formation of insoluble inclusions. Depletion of TDP-43 from the nucleus results in a loss of its normal function, which has emerged as a central driver of neurodegeneration^3,4^. The convergence of ALS and FTD on TDP-43 dysfunction suggests that understanding its normal roles in RNA processing – and how these are perturbed – could reveal fundamental mechanisms of disease and guide therapeutic strategies.

One of the key nuclear functions of TDP-43 is repression of cryptic splicing^5^. TDP-43 binds to specific intronic regions to suppress inclusion of these aberrant “cryptic” exons that would otherwise disrupt transcript integrity or even introduce new protein sequences. Beyond splicing, TDP-43 also regulates alternative polyadenylation, modulating 3′ UTR length and thereby influencing mRNA stability, localization, and translation^6–8^. The failure to repress cryptic exons or to maintain proper polyadenylation underscores TDP-43’s essential role in maintaining RNA homeostasis in the central nervous system.

Several cryptic splicing targets of TDP-43, including *STMN2*^9,10^, *UNC13A*^11,12^, and *KCNQ2*, have emerged as particularly compelling because their misprocessing contributes to neuronal dysfunction and degeneration, making them promising therapeutic entry points^13,14^. However, we hypothesized that these well-characterized targets may represent only the tip of the iceberg. Many cryptic exons introduced upon TDP-43 loss contain premature termination codons, resulting in rapid degradation of the mRNA via an mRNA surveillance mechanism called nonsense-mediated decay (NMD)^15^ (**Fig. 1a**). Case in point: *UNC13A*. The cryptic exon included in *UNC13A* upon TDP-43 loss introduces a premature termination codon, causing UPF1-dependent NMD^11^ and loss of *UNC13A* mRNA and protein^11,12^. This raises the possibility that RNA-sequencing approaches may underestimate the scope and the magnitude of cryptic splicing because transcripts subject to NMD could be depleted before detection. This might also explain why this *UNC13A* cryptic splicing event and other ones have shown variability in some patient lines^16^. To address this, we designed an experiment where we inhibited NMD along with TDP-43 knockdown prior to RNA-sequencing, aiming to stabilize and uncover otherwise hidden cryptic exon inclusion events that could reveal new pathogenic mechanisms and potential therapeutic targets.

**Figure 1.**
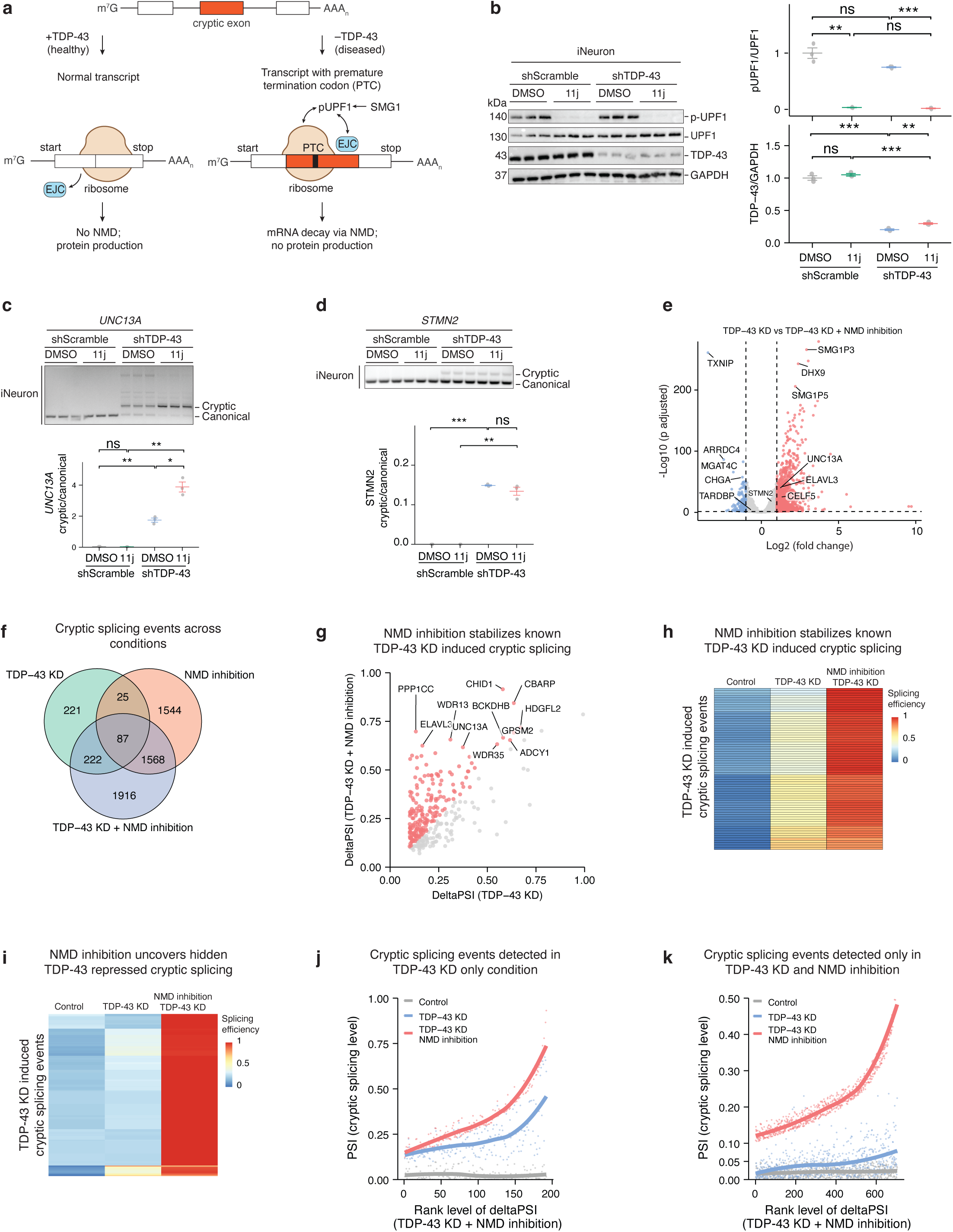
TDP-43-regulated cryptic splicing targets are substrates for nonsense-mediated decay. a. A schematic of nonsense-mediated decay (NMD) induced by cryptic splicing events. b. Western blots show that treating iNeurons with 0.5 µM of SMG1 inhibitor (11j) for 24 hours inhibited UPF1 phosphorylation and that compared to scramble shRNA virus (shScramble), TDP-43 shRNA virus (shTDP-43) efficiently knocked down TDP-43. c. RT-PCR shows that NMD inhibition by 11j treatment in iNeurons with TDP-43 knockdown (KD) increased the level of the cryptic isoform of *UNC13A*. d. RT-PCR shows that NMD inhibition by 11j did not increase the level of the cryptic isoform of *STMN2*. e. Volcano plot shows that combined NMD inhibition and TDP-43 knockdown in iNeurons caused widespread gene expression changes compared to TDP-43 knockdown alone. Genes with statistically significant changes (*p* adjusted < 0.05 and |log2(fold change)| >1) are labeled in red for increase and in blue for decrease. Dashed horizontal line marks the adjusted *p* value of 0.05 and vertical lines mark |log_2_(fold change)| of 1. f. Venn diagram shows the overlap of cryptic splicing events across three different conditions: TDP-43 knockdown, NMD inhibition by 11j treatment, and the combined NMD inhibition and TDP-43 knockdown. All conditions were compared to the control condition, where cells were treated with scramble shRNA and DMSO. g. Scatter plot shows NMD inhibition in TDP-43 knockdown increased cryptic splicing levels of known TDP-43 regulated cryptic splicing events. The deltaPSI in the x-axis represents the difference in cryptic splicing level (*PSI*) between TDP-43 knockdown and control (scramble shRNA + DMSO); the deltaPSI in the y-axis represents the difference in cryptic splicing level (*PSI*) between combined NMD inhibition and TDP-43 knockdown versus control (scramble shRNA + DMSO). Cryptic splicing levels (PSI) were calculated using leafcutter. h. Heatmap shows that for NMD sensitive, known TDP-43 cryptic splicing targets, their cryptic splicing levels (PSI) increased from control (scramble shRNA + DMSO) to TDP-43 knockdown to combined NMD inhibition by the 11j treatment and TDP-43 knockdown. i. Heatmap shows that for novel TDP-43 cryptic splicing targets, only combined NMD inhibition by 11j treatment and TDP-43 knockdown increased their cryptic splicing levels (PSI). j. Scatter plot shows that for NMD-sensitive, known TDP-43 cryptic splicing targets, we observed significant levels of cryptic splicing in both TDP-43 knockdown alone and in the combined NMD inhibition and TDP-43 knockdown condition. The solid line represents the linear model fit of each condition. k. Scatter plot shows that for NMD-sensitive, novel TDP-43 cryptic splicing targets, we only observed significant levels of cryptic splicing in the combined NMD inhibition and TDP-43 knockdown condition. The solid line represents the linear model fit of each condition. Unless otherwise stated, quantitation (*n* = 3 per condition) is presented as mean values +/- SEM; *p* values were two-sided Welch’s t-test; ns (not significant), *p* > 0.05; *, *p* ≤ 0.05; **, *p* ≤ 0.01; ***, *p* ≤ 0.001; ****, *p* ≤ 0.0001.

To test the hypothesis that TDP-43-dependent cryptic exon-containing mRNA transcripts are degraded by NMD, we knocked down TDP-43 in cortical neurons differentiated from human stem cells (iNeurons) and inhibited NMD using the pharmacological inhibitor 11j (**Fig. 1b**). NMD initiates, following the pioneering round of translation, when a complex of “up-frameshift proteins”, including UPF1-3, assembles on a prematurely terminating RNA^15,17^ (**Fig. 1a**). The kinase SMG1 phosphorylates the core member of the complex UPF1, which leads to the recruitment of enzymes and other factors that stimulate mRNA decay^18–20^. Since UPF1 phosphorylation is the rate-limiting step of NMD, the small molecule SMG1 inhibitor 11j has been used as a potent and selective NMD inhibitor^21^. Inhibting NMD with 11j for only 24 hours stabilized the *UNC13A* cryptic isoform in both TDP-43 knocked down iNeurons (**Fig. 1c**) and motor neurons derived from human induced pluripotent cells (iMNs) (**Fig. S1a,b**). Cryptic splicing in *STMN2* produces a truncated, polyadenylated mRNA that is not an NMD substrate^9,10^. Accordingly, NMD inhibition did not stabilize the *STMN2* cryptic isoform in either iNeurons (**Fig. 1d**) or iMNs (**Fig. S1c**).

To define the extent of TDP-43-dependent cryptic splicing events masked by NMD, we performed RNA-sequencing on iNeurons with or without TDP-43 knockdown and in the presence or absence of 11j. Compared to control samples, TDP-43 knockdown by itself (**Fig. S1d**), NMD inhibition by itself (**Fig. S1e**), or combined TDP-43 knockdown and NMD inhibition increased the levels of many transcripts (**Fig. 1e; S1f**), including ones harboring known TDP-43 repressed cryptic exons, such as *UNC13A*. Compared to TDP-43 knockdown alone, combined NMD inhibition and TDP-43 knockdown revealed 3,037 cryptic splicing events in 2,178 genes (*FDR* < 0.05 and *deltaPSI* > 0.1; **Fig. 1f**), of which 42% of these events (1,261) harbor TDP-43 binding sites within 500-nt. NMD inhibition in the TDP-43 knockdown background further increased splicing rates of 192 cryptic events detected in the TDP-43 knockdown alone condition, indicating that these TDP-43 dependent cryptic splicing events result in transcripts targeted for NMD (**Fig. 1g,h**).

In addition to cryptic splicing events detected by TDP-43 knockdown alone and by previous studies^11,12,22,23^, we reasoned that we might be able to detect cryptic events that were previously undetectable because of their rapid degradation by NMD. Combined NMD inhibition and TDP-43 knockdown unmasked 2,565 novel cryptic splicing events in 2,163 introns of 1,869 genes (*FDR* < 0.05 and *deltaPSI* > 0.1; **Fig. 1i**; **Fig. S1g**). Of these novel cryptic splicing events, 40% of them (1,029) harbor TDP-43 binding sites within 500-nt, 31% of events (795) showed at least 20% increase in splicing rate (*deltaPSI* > 0.2) compared to TDP-43 knockdown alone. Notably, TDP-43 knockdown alone detected splicing events showed high splicing rates in both TDP-43 knockdown and combined NMD inhibition and TDP-43 knockdown conditions (**Fig. 1j**), but cryptic splicing events that only became apparent upon NMD inhibition had minimal splicing in the TDP-43 knockdown alone condition (**Fig. 1k**), evidence of rapid transcript turnover by NMD. Consistent with this finding, a substantial fraction of genes (32%) harboring novel cryptic splicing events were downregulated upon TDP-43 knockdown (**Fig. S1h**). Together, these observations provide evidence that the vast majority of TDP-43 regulated cryptic splicing events lead to rapid transcript degradation by NMD.

NMD inhibition stabilized many known TDP-43-regulated cryptic splicing targets that are important for neuronal functions or are associated with diseases, such as *CAMK2B*, *KALRN, CDK7, ATG4B, MADD*, and *CELF5* (**Fig. 2a**). For example, we confirmed by RT-PCR in both iNeurons and iMNs (**Fig. 2b**) that NMD inhibition in the TDP-43 knockdown background further increased the cryptic isoform levels of *CELF5*, a gene encoding an RNA-binding protein that is highly expressed in the brain and involved in alternative splicing. Intriguingly, CELF5 protein was the third most downregulated protein in a recent mass spectrometry study of TDP-43 knockdown in iNeurons^23^. Notably, TDP-43 knockdown promoted cryptic splicing of *ELAVL3* (**Fig. 2c**), a gene that encodes a neuron-specific RNA-binding protein found to be one of the most downregulated in motor neurons of ALS patients and that harbors a cryptic exon 4a, whose inclusion results in an NMD-sensitive transcript^24,25^. NMD inhibition markedly stabilized the exon 4a containing cryptic isoform of *ELAVL3* (**Fig. 2c**), which we confirmed by RT-PCR in both iNeurons and iMNs (**Fig. 2d**). TDP-43 knockdown alone resulted in decreased full-length ELAVL3 protein level (**Fig. 2e**), providing further evidence that TDP-43 loss leads to cryptic exon inclusion and downregulation of *ELAVL3* mRNA transcripts (and resulting protein) via NMD.

**Figure 2.**
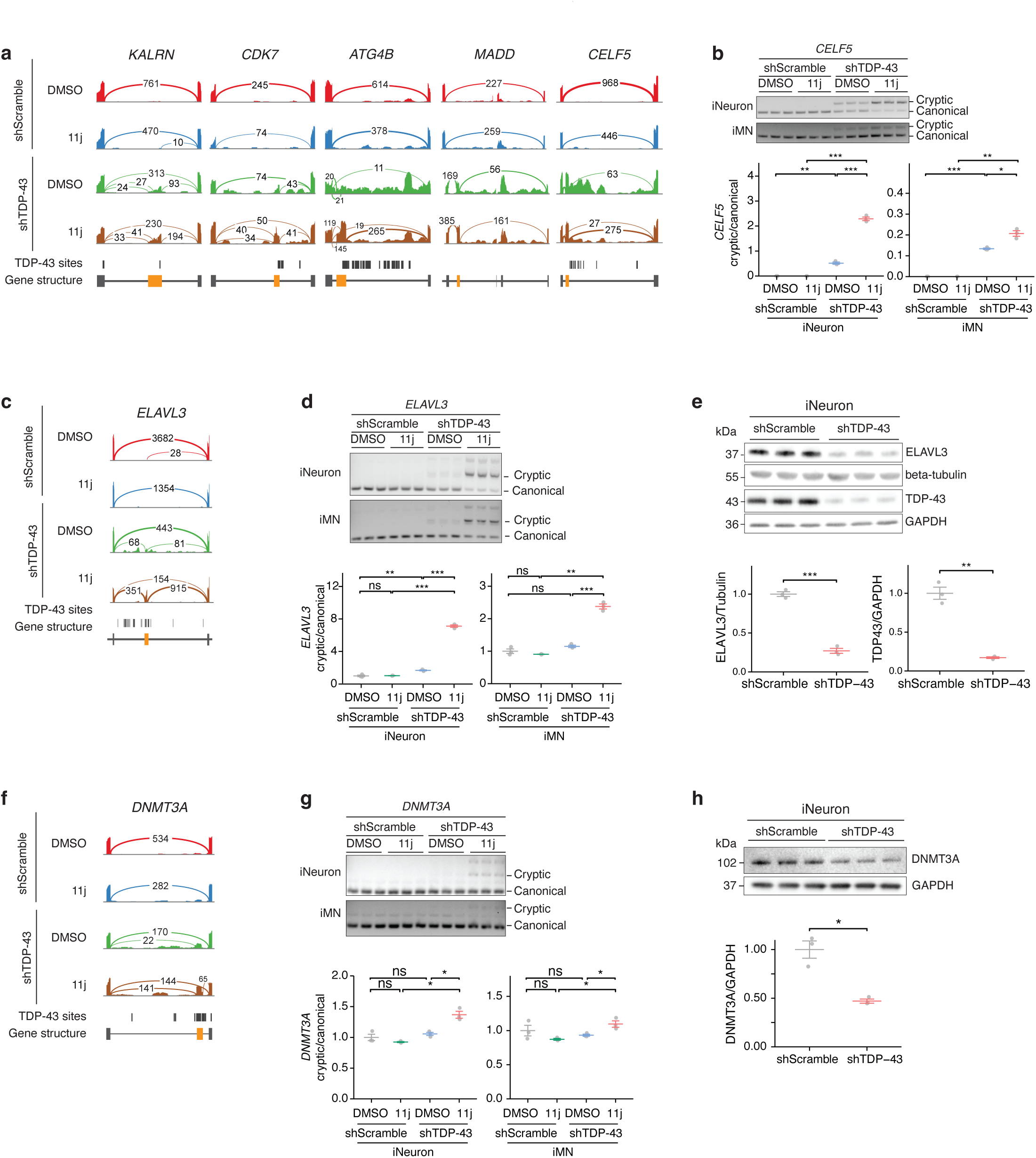
Inhibiting nonsense-mediated decay unmasks novel TDP-43-regulated cryptic splicing targets. a. Sashimi plots show examples of TDP-43 regulated cryptic splicing events across different conditions. The “TDP-43 sites” track marks the observed TDP-43 binding^33^. Gene structure is shown at the bottom. b. RT-PCR confirms that NMD inhibition by 11j treatment in iNeurons (top panel) or iMNs (bottom panel) with TDP-43 knockdown increased the level of the *CELF5* cryptic isoform. c. Sashimi plots show cryptic splicing in *ELALV3* across conditions. Details, as in **a**. d. RT-PCR confirms that NMD inhibition by 11j treatment in the TDP-43 knockdown background increased level of the cryptic isoform of *ELALV3* in iNeurons (top panel) and iMNs (bottom panel). e. Western blots show that TDP-43 knockdown reduced full-length ELAVL3 protein levels in iNeurons. f. Sashimi plots show cryptic splicing in *DNMT3A* across conditions. Details, as in **a**. g. RT-PCR confirms that only combined NMD inhibition by11j treatment and TDP-43 knockdown unmasked the cryptic isoform of *DNMT3A* in iNeurons (top panel) and iMNs (bottom panel). h. Western blots show that TDP-43 knockdown reduced full-length DNMT3A protein levels in iNeurons. Unless otherwise stated, quantitation (*n* = 3 per condition) is presented as mean values +/- SEM; *p* values were two-sided Welch’s t-test; ns (not significant), *p* > 0.05; *, *p* ≤ 0.05; **, *p* ≤ 0.01; ***, *p* ≤ 0.001; ****, *p* ≤ 0.0001.

NMD inhibition also unmasked many previously undetected TDP-43-regulated cryptic splicing targets (**Fig. 1e; Table S10**). For example, inhibiting NMD unmasked TDP-43-dependent cryptic exon usage in *DNMT3A* (**Fig. 2f**), a gene essential for DNA methylation^26,27^. We confirmed this by RT-PCR in both iNeurons and iMNs (**Fig. 2g**). Accordingly, TDP-43 knockdown reduced full-length DNMT3A protein levels (**Fig. 2h**). Loss of Dnmt3a in mouse causes motor neuron degeneration and decreased lifespan^28^, underscoring the importance of this gene for motor neuron function and suggesting a potential link between TDP-43 loss of function and DNA methylation abnormalities in neurons^29^. *DNMT3A* is one exemplar of how TDP-43-dependent cryptic splicing events that lead to a down regulation of protein levels can sometimes be masked and only become apparent upon NMD inhibition.

Here, we found that many (perhaps most) cryptic splicing events caused by loss of TDP-43 are rapidly degraded by NMD and undetectable by standard RNA-sequencing. By blocking NMD, we unmasked these cryptic splicing events. If most TDP-43 cryptic splicing events lead to NMD, how were we able to identify them in the first place^9–12^? Emerging evidence shows that TDP-43 loss can partially inhibit UPF1-dependent NMD^30,31^, potentially explaining why some of these have been difficult to detect in patient-derived neurons^16^. Therapeutic strategies aimed at targeting individual cryptic splicing events are being pursued^13,14^, but which ones are the right targets? Our results and similar ones by Sinha and colleagues^32^ provide evidence that there are many more TDP-43-dependent cryptic splicing events in neurons, which have previously been missed. Survivorship bias is the logical error of focusing only on things that pass a certain selection criteria while overlooking those that did not. This is illustrated by World War II analysts recommending armor only on places where returning planes were hit, not realizing the fatal hits were on non-returning aircraft. Here, by prioritizing cryptic splicing targets detected by RNA-sequencing, this bias ignores non-detected transcripts that were degraded by NMD. Thus, in addition to designing RNA-sequencing studies to account for NMD, mass spectrometry datasets^23^ will provide a valuable companion to assess the downstream impact of TDP-43 loss.

## Acknowledgments

We thank Derek Park and Neil Bhuteja for help with RT-PCR. We acknowledge funding support from the following sources: a postdoctoral scholar award from the Phil and Penny Knight Initiative for Brain Resilience, Stanford University (Y.Z.); a grant from the Larry L. Hillblom foundation (Y.Z.); the National Institutes of Health (NIH) Training Program in Aging Research (2T32AG047126-06A1 to T.A.); a grant from the Takeda Science Foundation (T.A.); the NIH (U54NS123743, R35NS097273, P01NS084974) and Target ALS to L.P.; and the NIH (R35NS137159, U54NS123743, R01AG064690) and Target ALS to A.D.G.. A.D.G. is a Chan Zuckerberg Biohub – San Francisco Investigator. Some of the computing for this project was performed on the Sherlock cluster. We would like to thank Stanford University and the Stanford Research Computing Center for providing computational resources and support that contributed to these research results.

## Methods

### Stem cell maintenance and differentiation into iNeurons and iMNs

Human embryonic stem cells (hESCs; H1) were purchased from WiCell (WA01), authenticated by WiCell. Induced pluripotent stem cells (iPSCs; WTC11) were purchased from Coriell Institute for Medical Research (GM25256). H1 and WTC11 cells were maintained in mTeSR Plus media (StemCell Technologies, 100-0276) with 50 U/mL Penicillin-Streptomycin (Gibco, 15140122) on plates coated with Matrigel (Corning, 354230). Cells were fed every two days and split every 3-4 days using ReLeSR (StemCell Technologies, 100-0483) according to the manufacturer’s instructions. WTC11 cells were split using mTeSR Plus with 10nM of ROCK1 inhibitor (Tocris, 1254), and the media was changed to one without ROCK1 inhibitor within 24 hours. The differentiation of hESCs to neurons by forcing NGN2 over-expression was carried out as previously described^34^. In brief, cells were transduced with a Tet-On induction system to drive the expression of the transcription factor NGN2.

Cells were dissociated on day 3 of differentiation and replated on poly-D-Lysine and Laminin double-coated tissue culture plates in Neurobasal Medium (Thermo Fisher, 21103049) containing neurotrophic factors, BDNF (Peprotech 450-10) and GDNF (Peprotech 450-03). The differentiation of WTC11 iPSCs to iMNs was conducted by forcing the over expression of NGN2, ISL1, and LHX3 using the Tet-On induction system. The cells were replated on day 3 on poly-D-Lysine and Laminin double-coated tissue culture plates in DMEM/F12 containing ROCK inhibitor, doxycycline (Sigma Aldrich, D9891-5G), Compound E (Abcam, ab142164), laminin (Sigma Aldrich, L2020), and BrdU (Millipore Sigma, B9285). On day 4, media was changed into DMEM/F12 (ThermoFisher Scientific, 11330032) containing neurotrophic factors, BDNF, GDNF, and NT3 (Peprotech, 450-03), laminin, and polB inhibitor Aphidicolin (Millipore Sigma, 89458). On day 10, media was changed into maintenance media using Neurobasal Medium containing neurotrophic factors, laminin and Aphidicolin.

### TDP-43 knockdown in iNeurons and iMNs

After seven days of being cultured for differentiation, iNeurons and iMNs were transduced with lentivirus expressing scrambled shRNA or TDP-43 shRNA as previously described^12^, cultured for additional eight days, and then collected for downstream analyses. The knockdown efficiency was assessed by Western blotting.

### Inhibition of the nonsense-mediated decay

After treating scramble shRNA or TDP-43 shRNA virus for seven days, iNeurons and iMNs were treated with 0.5uM of 11j (MedChem Express, HY-124719), a SMG-1 inhibitor, or an equal volume of DMSO for 24 hours. The concentration of 11j was determined in iNeurons previously^8^. Cells were harvested for RNA and protein lysates. Western blot was used to confirm that phosphorylation of UPF1 was inhibited. Western blot was performed as described below with wet transfer and proteins were detected using the following antibodies: UPF1 (1:10,000; Abcam ab109363), pUPF1 (1:1,000; EMD Millipore, 07-1016), and GAPDH (1:2,000; Sigma-Aldrich, G8795).

### Extraction of total RNA and proteins from iNeurons and iMNs

Total RNA and protein were extracted using AllPrep DNA/RNA/Protein Mini Kit (Qiagen, 80004) according to the manufacturer’s instructions, except that protein pellets were resuspended in 60 uL of 5% SDS solution.

### Immunoblotting

The protein lysates were used for bicinchoninic acid (Invitrogen, 23225) assays to determine protein concentrations. Unless stated otherwise, 5-10 µg of protein lysates from each sample was denatured for 10 min at 85 °C in Laemmli sample buffer (Bio-Rad, 1610747), containing 2-mercaptoethanol (Sigma-Aldrich). These samples were loaded onto 4–12% Bis–Tris Mini gels (Thermo Fisher, NP0336BOX) for gel electrophoresis and then transferred onto 0.45-μm nitrocellulose membranes (Bio-Rad, 162-0115) using the semi-dry transfer method (Bio-Rad Trans-Blot Turbo Transfer System, 1704150) or at 120 V for 2 h at 4 °C using the wet transfer method (Bio-Rad Mini Trans-Blot Electrophoretic Cell, 170-3930) onto PVDF membranes (Bio-Rad, 1620219). Membranes were blocked in EveryBlot Blocking Buffer (Bio-Rad, 12010020) for 1 h then incubated overnight at 4°C in blocking buffer containing antibodies against phosphorylated-UPF1 (1:1,000; EMD Millipore, 07-1016 ), UPF1(1:10,000; Abcam, ab109363), TDP-43 (1:1,500; Proteintech, 10782-2-AP), ELAVL3 (1:1,000; Proteintech, 55047-1-AP), DNMT3A(1:500; Abcam, ab2850), Beta Tubulin (1:40,000; Proteintech, 66240-1-Ig), GAPDH (1:2,000; Sigma-Aldrich, G8795), Histone H3 (1:5,000; Abcam, ab1791).

Membranes were subsequently incubated in blocking buffer containing horseradish peroxidase (HRP)- conjugated anti-mouse IgG (H+L) (1:5,000, Thermo Fisher, 62-6520) or HRP-conjugated anti-rabbit IgG (H+L) (1:5,000, Life Technologies, 31462) for 1 h. The Amersham ECL Prime kit (Cytiva, RPN2232) or SuperSignal™ West Femto Maximum Sensitivity Substrate (Thermo Fisher, 34094) was used to develop blots, which were then imaged using ChemiDox XRS+ System (Bio-Rad). The intensity of bands was quantified using Fiji and then normalized to the corresponding controls.

### RT-PCR

Total RNA (500 ng) was reverse transcribed to cDNA using the PrimeScript™ RT Reagent Kit with gDNA Eraser (Takara, RR047A). PCR was conducted using Q5 High Fidelity 2x Master mix (NEB, M0492). The resulting PCR products were visualized on a 2% agarose gel. Primers and expected PCR product sizes are listed in Table S11.

### Competition PCR for *STMN2*

Total RNA (500 ng) was reverse transcribed to cDNA using the PrimeScript™ RT Reagent Kit with gDNA Eraser (Takara, RR047A). Competition PCR was carried out using the Q5 High Fidelity 2x Master mix (NEB, M0492) using one forward primer and two reverse primers —one targeting exon 2 and the other targeting the cryptic exon. The resulting PCR products were visualized on a 2% agarose gel. Primers are listed in Table S11.

### Total RNA sequencing

Total RNA was used to construct RNA-sequencing libraries using the SMARTer Stranded Total RNA-Seq Kit v2 - Pico Input Mammalian kit (Takara, 634411), according to the manufacturer’s instructions. The resulting libraries were quantitated, pooled, and sequenced on a 10-billion flowcell on an Illumina Novaseq X machine.

### Gene expression analysis

Adapters in FASTQ files were trimmed using fastp. The adapter trimmed FASTQ files were used for differential gene expression analysis using Salmon and DESeq2.

### Splicing analysis

The adapter-trimmed FASTQ files were mapped to the human genome (hg38) following ENCODE’s recommended settings using STAR. The unique-mapping, properly paired reads were then used for splicing analysis using LeafCutter. Cryptic splicing events were called by LeafCutter. To identify novel cryptic splicing events, cryptic splicing events were filtered if they were detected in TPD-43 knockdown alone condition or reported previously.

### Quantitation and statistical analysis

All quantification and statistical analyses were done in R. Analysis details can be found in figure legends, methods, and the main text. Genome tracks were prepared using IGV. All plots were prepared using tidyplots, ggplot2, ggpubr, patchwork, and ggrepel in R.

## Data and code availability

The corresponding sequencing data generated in this study will be available at GEO. All processed data are listed in supplementary tables. No new code is used in this study. The following publicly available sequencing data are used in this study: GSE126542 and ERP126666.

## List of supplementary tables

Table S1: Gene expression analysis result of TDP-43 knockdown vs control by RNA-seq

Table S2: Gene expression analysis result of 11j treatment vs control by RNA-seq

Table S3: Gene expression analysis result of TDP-43 knockdown and 11j treatment vs control by RNA-seq

Table S4: Gene expression analysis result of TDP-43 knockdown and 11j treatment vs TDP-43 knockdown by RNA-seq

Table S5: Cryptic splicing events detected in TDP-43 knockdown vs control samples

Table S6: Cryptic splicing events detected in 11j treatment vs control samples

Table S7: Cryptic splicing events detected in TDP-43 knockdown and 11j treatment vs control samples

Table S8: Cryptic splicing events detected in TDP-43 knockdown and 11j treatment vs TDP-43 knockdown samples

Table S9: Cryptic splicing events detected in TDP-43 knockdown with increased rate in TDP-43 and 11j treatment

Table S10: Novel cryptic splicing events revealed by TDP-43 knockdown and 11j treatment Table S11: List of primers used in RT-PCR

---

**Figure S1.**
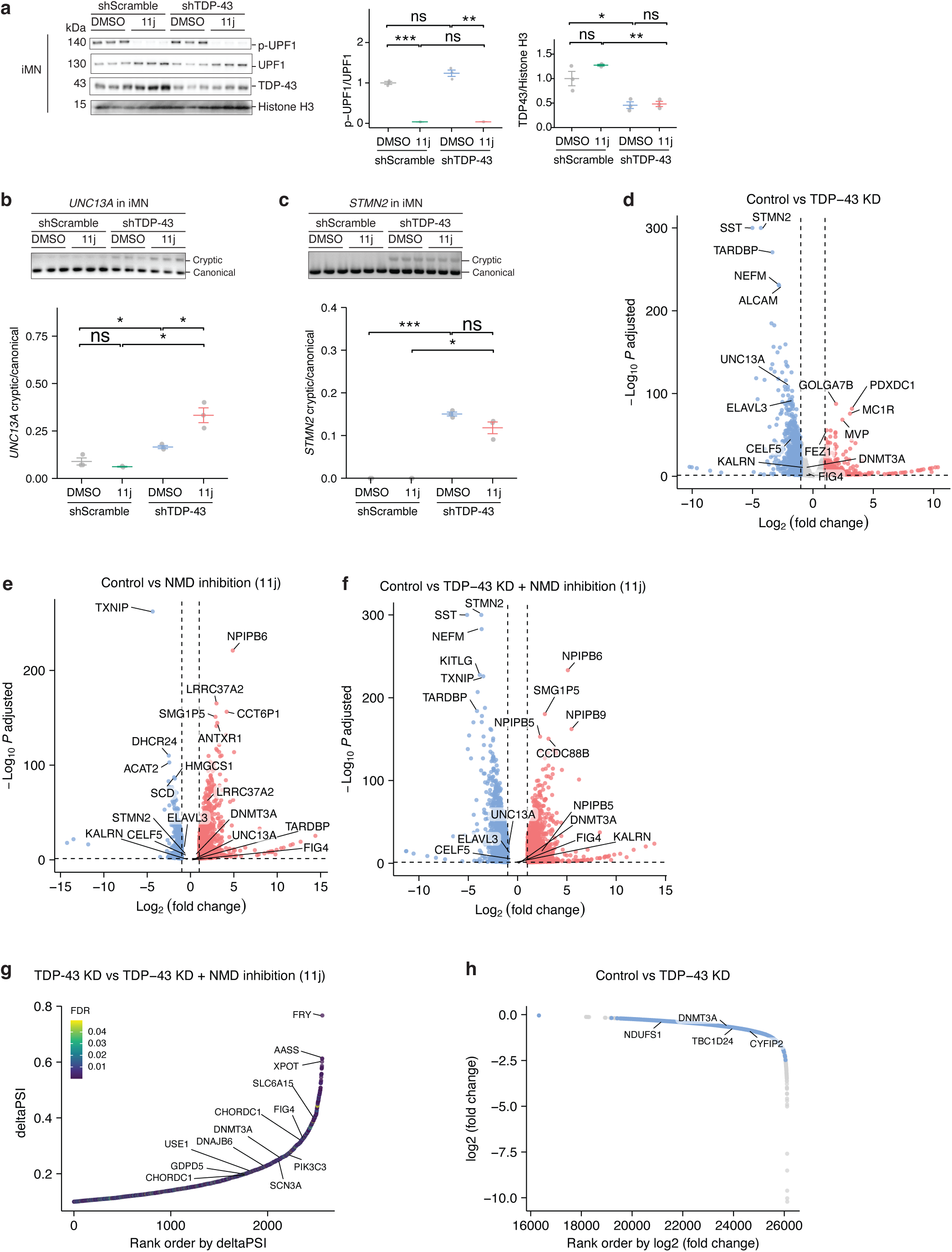
TDP-43 loss-induced cryptic splicing targets transcripts for nonsense mediated decay. a. Western blots show iMNs with and without TDP-43 knockdown (KD) treated with 0.5 µM of SMG1 inhibitor (11j) for 24 hours, which inhibited UPF1 phosphorylation. Details as in Fig. 1b. b. RT-PCR shows that nonsense-mediated decay (NMD) inhibition by 11j treatment in the TDP-43 knockdown background increased level of the cryptic isoform of *UNC13A* in iMNs. Details as in Fig. 1c. c. RT-PCR shows that NMD inhibition by 11j treatment in the TDP-43 knockdown background did not increase level of the cryptic isoform of *STMN2* in iMNs. Details as in Fig. 1c. d. Volcano plot shows that TDP-43 knockdown in iNeurons caused gene expression changes. Details, as in Fig. 1e. e. Volcano plot shows that NMD inhibition by 11j treatment in iNeurons caused gene expression changes. Details, as in Fig. 1e. f. Volcano plot shows that combined NMD inhibition by 11j treatment and TDP-43 knockdown in iNeurons caused gene expression changes compared to the control condition (scramble shRNA + DMSO). Details, as in Fig. 1e. g. Scatter plot shows cryptic splicing increase of novel TDP-43 cryptic splicing events (deltaPSI on the y-axis) ordered by their ranks. False discovery rate (FDR) was calculated using leafcutter. h. Scatter plot shows gene expression reduction upon TDP-43 knockdown on the y-axis (log2(fold change)) ordered by their gene expression ranks. Genes with novel cryptic splicing that showed expression reduction is labeled in blue.

## References

1. Taylor, J. P., Brown, R. H. & Cleveland, D. W. Decoding ALS: from genes to mechanism. Nature 539, 197–206 (2016).

2. Neumann, M. et al. Ubiquitinated TDP-43 in Frontotemporal Lobar Degeneration and Amyotrophic Lateral Sclerosis. Science 314, 130–133 (2006).

3. Klim, J. R., Pintacuda, G., Nash, L. A., Juan, I. G. S. & Eggan, K. Connecting TDP-43 Pathology with Neuropathy. Trends in Neurosciences 44, 424–440 (2021).

4. Kim, G., Gautier, O., Tassoni-Tsuchida, E., Ma, X. R. & Gitler, A. D. ALS Genetics: Gains, Losses, and Implications for Future Therapies. Neuron 108, 822–842 (2020).

5. Ling, J. P., Pletnikova, O., Troncoso, J. C. & Wong, P. C. TDP-43 repression of nonconserved cryptic exons is compromised in ALS-FTD. Science 349, 650–655 (2015).

6. Arnold, F. J. et al. TDP-43 dysregulation of polyadenylation site selection is a defining feature of RNA misprocessing in amyotrophic lateral sclerosis and frontotemporal dementia. J Clin Invest 135, (2025).

7. Bryce-Smith, S. et al. TDP-43 loss induces extensive cryptic polyadenylation in ALS/FTD. 2024.01.22.576625 Preprint at 10.1101/2024.01.22.576625 (2024).

8. Zeng, Y. et al. TDP-43 nuclear loss in FTD/ALS causes widespread alternative polyadenylation changes. 2024.01.22.575730 Preprint at 10.1101/2024.01.22.575730 (2024).

9. Klim, J. R. et al. ALS-implicated protein TDP-43 sustains levels of STMN2, a mediator of motor neuron growth and repair. Nat Neurosci 22, 167–179 (2019).

10. Melamed, Z. et al. Premature polyadenylation-mediated loss of stathmin-2 is a hallmark of TDP-43-dependent neurodegeneration. Nat Neurosci 22, 180–190 (2019).

11. Brown, A.-L. et al. TDP-43 loss and ALS-risk SNPs drive mis-splicing and depletion of UNC13A. Nature 1–8 (2022) doi:10.1038/s41586-022-04436-3.

12. Ma, X. R. et al. TDP-43 represses cryptic exon inclusion in the FTD–ALS gene UNC13A. Nature 603, 124–130 (2022).

13. Baughn, M. W. et al. Mechanism of STMN2 cryptic splice-polyadenylation and its correction for TDP-43 proteinopathies. Science 379, 1140–1149 (2023).

14. Keuss, M. J. et al. Loss of TDP-43 induces synaptic dysfunction that is rescued by UNC13A splice-switching ASOs. bioRxiv 2024.06.20.599684 (2024) doi:10.1101/2024.06.20.599684.

15. Kurosaki, T., Popp, M. W. & Maquat, L. E. Quality and quantity control of gene expression by nonsense-mediated mRNA decay. Nat Rev Mol Cell Biol 20, 406–420 (2019).

16. Rothstein, J. D., Warlick, C. & Coyne, A. N. Highly variable molecular signatures of TDP-43 loss of function are associated with nuclear pore complex injury in a population study of sporadic ALS patient iPSNs. *bioRxiv* 2023.12.12.571299 (2023) doi:10.1101/2023.12.12.571299.

17. Lykke-Andersen, S. & Jensen, T. H. Nonsense-mediated mRNA decay: an intricate machinery that shapes transcriptomes. Nat Rev Mol Cell Biol 16, 665–677 (2015).

18. Yamashita, A., Ohnishi, T., Kashima, I., Taya, Y. & Ohno, S. Human SMG-1, a novel phosphatidylinositol 3-kinase-related protein kinase, associates with components of the mRNA surveillance complex and is involved in the regulation of nonsense-mediated mRNA decay. Genes Dev 15, 2215–2228 (2001).

19. Conti, E. & Izaurralde, E. Nonsense-mediated mRNA decay: molecular insights and mechanistic variations across species. Curr Opin Cell Biol 17, 316–325 (2005).

20. Lejeune, F. & Maquat, L. E. Mechanistic links between nonsense-mediated mRNA decay and pre-mRNA splicing in mammalian cells. Curr Opin Cell Biol 17, 309–315 (2005).

21. Gopalsamy, A. et al. Identification of pyrimidine derivatives as hSMG-1 inhibitors. Bioorg Med Chem Lett 22, 6636–6641 (2012).

22. Liu, E. Y. et al. Loss of Nuclear TDP-43 Is Associated with Decondensation of LINE Retrotransposons. Cell Reports 27, 1409–1421.e6 (2019).

23. Seddighi, S. et al. Mis-spliced transcripts generate de novo proteins in TDP-43-related ALS/FTD. Sci Transl Med 16, eadg7162 (2024).

24. Diaz-Garcia, S. et al. Nuclear depletion of RNA-binding protein ELAVL3 (HuC) in sporadic and familial amyotrophic lateral sclerosis. Acta Neuropathol 142, 985–1001 (2021).

25. Costantino, I., Meng, A. & Ravits, J. Alternatively spliced ELAVL3 cryptic exon 4a causes ELAVL3 downregulation in ALS TDP-43 proteinopathy. Acta Neuropathol 147, 93 (2024).

26. Okano, M., Xie, S. & Li, E. Cloning and characterization of a family of novel mammalian DNA (cytosine-5) methyltransferases. Nat Genet 19, 219–220 (1998).

27. Okano, M., Bell, D. W., Haber, D. A. & Li, E. DNA methyltransferases Dnmt3a and Dnmt3b are essential for de novo methylation and mammalian development. Cell 99, 247–257 (1999).

28. Nguyen, S., Meletis, K., Fu, D., Jhaveri, S. & Jaenisch, R. Ablation of de novo DNA methyltransferase Dnmt3a in the nervous system leads to neuromuscular defects and shortened lifespan. Dev Dyn 236, 1663–1676 (2007).

29. Hop, P. J. et al. Genome-wide study of DNA methylation shows alterations in metabolic, inflammatory, and cholesterol pathways in ALS. Sci Transl Med 14, eabj0264 (2022).

30. Gomez, N. et al. Counter-regulation of RNA stability by UPF1 and TDP43. 2024.01.31.578310 Preprint at 10.1101/2024.01.31.578310 (2024).

31. Alessandrini, F., Wright, M., Kurosaki, T., Maquat, L. E. & Kiskinis, E. ALS-Associated TDP-43 Dysfunction Compromises UPF1-Dependent mRNA Metabolism Pathways Including Alternative Polyadenylation and 3’UTR Length. 2024.01.31.578311 Preprint at 10.1101/2024.01.31.578311 (2024).

32. Sinha, I. R. et al. Inhibition of nonsense-mediated decay in TDP-43 deficient neurons reveals novel cryptic exons. 2025.06.28.661837 Preprint at 10.1101/2025.06.28.661837 (2025).

33. Zhao, W. et al. POSTAR3: an updated platform for exploring post-transcriptional regulation coordinated by RNA-binding proteins. Nucleic Acids Res 50, D287–D294 (2022).

34. Bieri, G. et al. LRRK2 modifies α-syn pathology and spread in mouse models and human neurons. Acta Neuropathol 137, 961–980 (2019).

